# Epigenetic MRI: Noninvasive Imaging of DNA Methylation in the Brain

**DOI:** 10.1101/2021.08.20.457113

**Authors:** Fan Lam, James Chu, Ji Sun Choi, Chang Cao, T. Kevin Hitchens, Scott K. Silverman, Zhi-Pei Liang, Ryan N. Dilger, Gene E. Robinson, King C. Li

## Abstract

Both neuronal and genetic mechanisms regulate brain function. While there are excellent methods to study neuronal activity in vivo, there are no nondestructive methods to measure global gene expression in living brains. Here we present a method, epigenetic magnetic resonance imaging (eMRI), that overcomes this limitation via direct imaging of DNA methylation, a major gene expression regulator. eMRI exploits the methionine metabolic pathways for DNA methylation to label genomic DNA through ^13^C-enriched diets. A novel ^13^C magnetic resonance spectroscopic imaging method then maps the spatial distribution of labeled DNA. We validated eMRI using pigs, whose brains have stronger similarity to humans in volume and anatomy than rodents, and confirmed efficient ^13^C labeling of brain DNA. We also discovered strong regional differences in global DNA methylation. Just as MRI measurements of regional neuronal activity have had a transformational effect on neuroscience, we expect that the eMRI signal as a surrogate for regional gene expression will enable many new investigations into the roles of gene expression in human brain function, behavior, and disease.

## Introduction

The brain is ever-changing in structure and function as a result of development, aging, environmental influence, and disease. Two fundamental mechanisms underpin these changes: neuronal activation, which occurs over relatively short time scales (milliseconds, seconds, and minutes), and gene expression, which occurs over longer time scales (hours, days, or even longer) ^1–3^. Advances in imaging technology have transformed how we investigate these mechanisms.

Functional magnetic resonance imaging (fMRI) has revolutionized our understanding of the human brain by providing a powerful nondestructive method to image neural activity ^4–7^. In contrast, the technologies to image gene expression have been limited to methods that require invasive sampling and tissue processing ^8–11^. Although these techniques have provided tremendous knowledge about gene expression and gene regulation in the brain, especially in animal models, their destructive nature makes longitudinal studies of the same samples impossible, thus limiting our ability to translate and expand scientific discoveries to human brains. This is especially unfortunate because longer-term changes in brain function play critical roles in both brain diseases and responses of the brain to environmental change ^1–3^. The ability to measure and visualize gene expression and regulation in the brain noninvasively would revolutionize the study of brain function, behavior, and disease ^12^.

Efforts to map brain gene expression and regulation in living organisms to date have involved imaging reporter genes or associated enzymes using optical techniques, PET, or MRI ^13–17^. These methods are either limited to model organisms or require transgenic animals engineered to express a particular reporter gene and an exogenous contrast probe interacting with the reporter gene to produce the desired images ^15,16^. Therefore, such methods have no clear path for translation to humans. Furthermore, these methods are limited to just a few genes and therefore cannot provide a comprehensive portrait of gene expression.

PET imaging of histone deacetylases (HDACs) in the human brain was recently demonstrated, using a radioactive tracer that can pass the blood-brain barrier (BBB) and target HDAC isoforms ^18,19^. This probes a major form of epigenetic gene regulation, histone acetylation, but PET requires introducing radioactive materials into the body, and it only targets one of the enzymes that regulates histone acetylation rather than histone acetylation itself. Moreover, PET lacks the specificity to distinguish between the target molecule and downstream metabolic products ^20–22^.

Other imaging epigenetics approaches have studied correlations between brain MRI and gene expression or methylation in postmortem tissues, saliva or blood ^23–28^; although useful, they can only provide indirect insights into the brain gene expression and regulation.

We present the first successful direct imaging of brain DNA methylation using a novel approach we call epigenetic MRI (eMRI), which integrates stable isotope 5-methyl-2′-deoxycytidine (5mdC) labeling through diet and magnetic resonance spectroscopic imaging (MRSI). Using pigs fed by a customized diet enriched in ^13^C-methionine (^13^C-Met) and innovations to ^13^C-MRSI, we report robust mapping in intact brain hemispheres that revealed strong regional differences in DNA methylation. We chose pigs as a surrogate to assess feasibility of translation to humans because of stronger similarities in brain size and anatomy than rodent models ^29^. Significant eMRI signal differences were observed in animals fed with enriched diet for different numbers of days, demonstrating the dynamic nature of this signal. Given the noninvasiveness of our method, these results provide a path towards a global DNA methylation brain imaging paradigm for humans. Because DNA methylation is one of the major regulators of gene expression, eMRI promises to become a powerful tool to understand the molecular basis of brain function and disease.

## Results

### In vivo brain DNA methylation labeling using ^13^C-methionine-enriched diet

A key feature of eMRI is the use of a natural amino acid involved in DNA methylation that is labeled with an NMR-active stable isotope, making methylated DNA detectable with noninvasive imaging (Fig. 1). We chose ^13^C-Met because Met is an essential amino acid and the major methyl group donor for DNA methylation. Met is also commonly used as a nutritional supplement and approved for human use. To validate our labeling approach, we designed a special diet that replaced all protein with free amino acids in the proper proportions and substituted all Met with enriched ^13^C-Met.

**Fig. 1.**
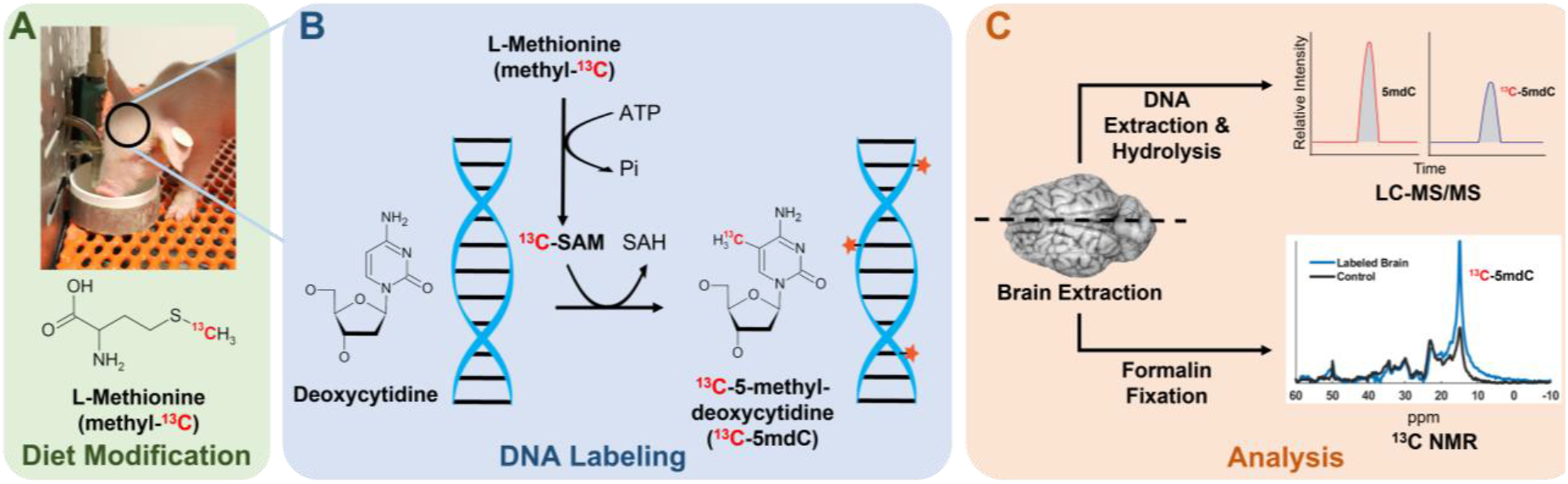
Design of dietary ^13^C labeling of genomic DNA in the brain and eMRI. **A**, A special diet enriched with ^13^C-Met was ingested by neonatal piglets through a milk replacement formula for either 10 or 32 days. Diet for age-matched controls had all ^13^C-Met replaced by regular Met. **B**, Ingested ^13^C-Met passes through the BBB and is converted to ^13^C-SAM, which methylates the DNA, producing ^13^C-methyl-labeled DNA. **C**, Brain tissues were sampled from one hemisphere for DNA extraction and LC-MS/MS analysis to validate labeling. The other intact hemisphere was analyzed using nondestructive ^13^C NMR and MRSI, another key feature of eMRI.

We fed this diet to piglets starting from postnatal day 1 (Fig. 1A). The ingested ^13^C-Met crosses the BBB ^30–32^ and is converted into ^13^C-*S*-adenosylmethionine (^13^C-SAM), which then provides the ^13^C-methyl group for cytosine methylation, effectively labeling the 5mdC in brain genomic DNA (Fig. 1B). This is similar to the labeling scheme used to study the conversion of 5mdC to 5hmdC (5-hydroxymethyl-2′-deoxycytidine) in laboratory mice with triply deuterated ^13^C-Met ^33^. After diet replacement for 10 or 32 days, animals were sacrificed. Following brain dissection, one hemisphere was used for tissue sampling from various brain regions, DNA extraction, and analysis by LC-MS/MS to confirm DNA labeling; the other intact hemisphere was used for noninvasive eMRI (Fig. 1C). Control animals were fed with the same formula for 10 or 32 days, but all ^13^C-Met was replaced with non-isotope-labeled Met.

LC-MS/MS analysis revealed effective labeling of brain 5mdC after feeding for 10 days with the ^13^C-Met-enriched diet. Relative to controls, we detected a 3–4% increase in ^13^C-labeled 5mdC (as a fraction of total 5mdC) across eight different sampled brain regions (Fig. 2A). Feeding for 32 days resulted in much stronger labeling, with >12% increase in ^13^C-5mdC compared to the age-matched controls (Fig. 2B). The baseline ^13^C-labeled 5mdC signal was ∼5% in the controls at both ages (Fig. 2C), which is consistent with the 1.1% natural abundance of the ^13^C isotope and accounting for the five carbons in 5mdC ^34^.

**Fig. 2.**
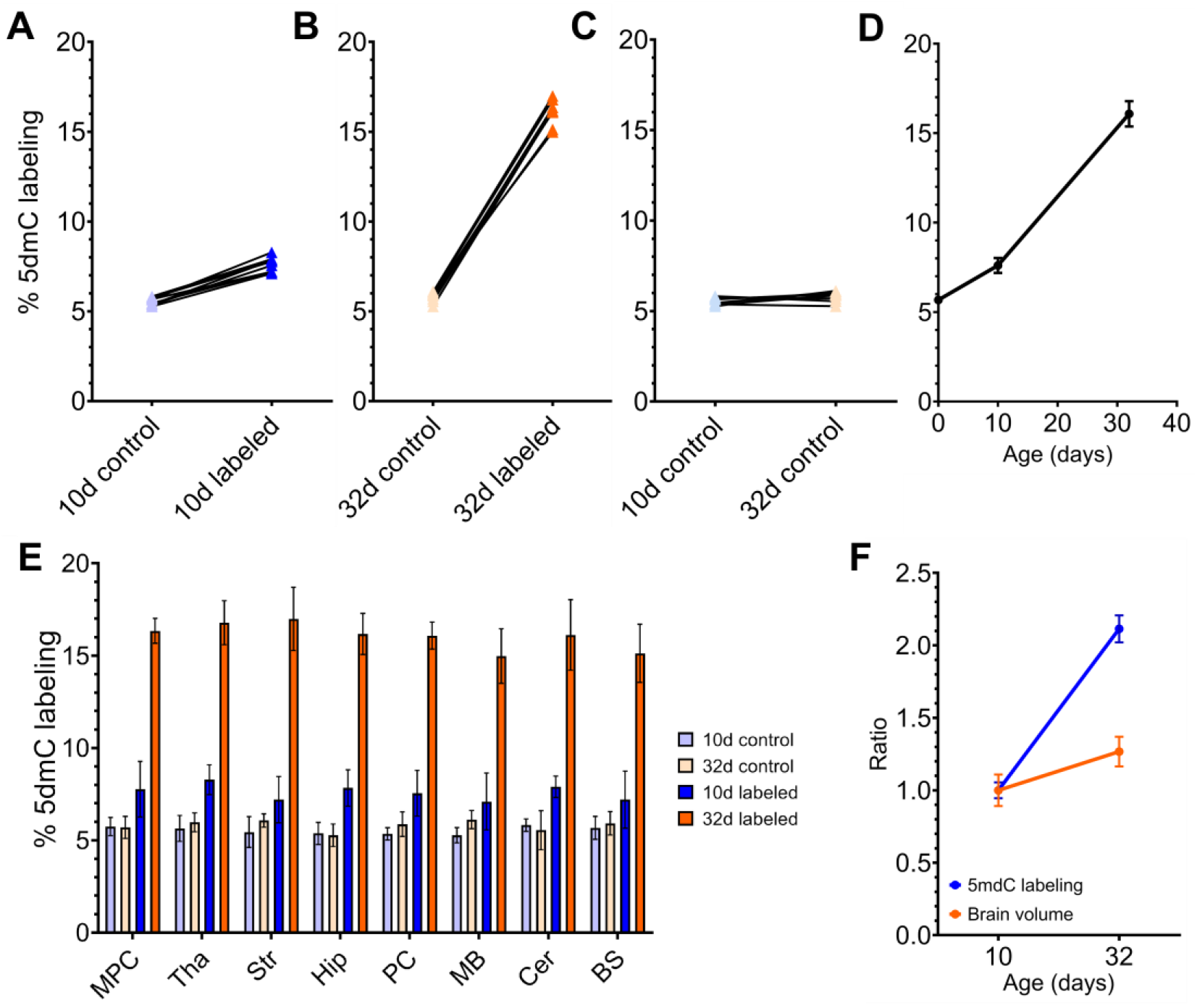
Dietary ^13^C labeling of brain genomic DNA confirmed by LC-MS/MS. **A**, After 10 days of the ^13^C-Met-enriched diet, ^13^C-5mdC in the brain DNA (ratio of ^13^C-5mdC to total 5mdC) increased from ∼5% to 8–9%. **B**, After 32 days ^13^C-5mdC in the brain DNA significantly increased to ∼15–18%. **C**, Age-matched controls for both 10-day and 32-day diets had ∼5% ^13^C-5mdC, consistent with the 1.1% natural abundance of the ^13^C isotope and validating our LC-MS/MS analysis. **D**, Because ∼5% ^13^C-5mdC can be assumed at birth, we back-extrapolated the curve for percentage versus days of diet. A nonlinear increase in labeling was observed, with the increase from day 10 to 32 much greater than for day 0 to 10. **E**, A more detailed comparison of labeling percentages for different diet groups and brain regions; error bars indicate standard deviations. No regional differences were observed for the controls. **F**, The ^13^C-5mdC increase from 10 to 32 days with ^13^C diet (∼2×) is considerably larger than the brain volume increase that occurred over the same time period (∼20%). Our calibration curve with MS peak areas versus concentrations of 5mdC confirms that the range reported here is within the linear range of our LC-MS/MS protocol (data not shown). Brain regions: medial prefrontal cortex (MPC), thalamus (Tha), striatum (Str), hippocampus (Hip), perirhinal cortex (PC), midbrain (MB), cerebellum (Cer), brain stem (BS).

The 32-day labeled samples produced >2× labeling percentage compared to the 10-day labeled samples (Fig. 2D, P < 0.001, one-way ANOVA). As expected, the baseline ^13^C-5mdC percentage did not differ among the eight brain regions or between the control animals fed for either 10 or 32 days (Fig. 2C, E). Differences in ^13^C-5mdC percentage across brain regions in the labeled samples were not statistically significant for either time point (Fig. 2E). Representative LC-MS/MS data revealing the ^13^C-5mdC and 5mdC peaks are in Supplementary Fig. 1.

From these data, two interesting observations were made about DNA methylation dynamics in the brain. First, there was a nonlinear increase in labeling from day 10 to 32, much greater than from day 0 to 10 (Fig. 2D). This increase occurred despite a relative decrease in daily ^13^C-Met intake (in terms of mg/kg body weight/day) due to rapid increases in body weight over the same time period. We speculate this to be the result of an increase in DNA methylation events and the methylation-demethylation turnover cycle as part of normal brain development. Second, the ∼2-fold increase in labeling percentage detected in animals fed ^13^C-Met for 32 days relative to 10 days vastly outpaced the ∼20% increase in brain volume that occurred during the same period (Fig. 2F). This finding suggests that increases in tissue volume and neuronal density during brain development are not the only drivers of increased ^13^C-methyl incorporation into brain DNA. This implicates either intrinsic age-related changes in methylation dynamics or extrinsic processes such as learning and memory formation, which are known to affect brain DNA methylation ^1– 3,35–37^. They may cause unlabeled methyl groups to be replaced with ^13^C-labeled methyl groups through turnover or cause unmethylated cytosine to be methylated. These results also demonstrate the responsiveness of the eMRI signal to changing experimental conditions.

### Noninvasive detection of labeled methylated DNA using ^13^C NMR

Strong isotope labeling of methylated DNA in the brain enables the possibility of using noninvasive isotopic detection methods. For eMRI, we chose ^13^C NMR to detect and quantify the ^13^C-labeled 5mdC in genomic DNA. ^13^C NMR and imaging using isotope-enriched molecules has been used in various basic biological studies and clinical applications ^38–41^. It offers several unique advantages: 1) Its nondestructive and nonradioactive nature allows for in vivo translation to humans; 2) Its broad chemical shift dispersion allows for detecting signals specifically from ^13^C-5mdC in genomic DNA relative to ^13^C signals from other molecules; and 3) ^13^C has low natural abundance (∼1.1%), which can increase specific signal detection from the ^13^C-enriched diet.

We first performed a proof-of-principle in vitro experiment using synthetic DNA oligonucleotides with a well-defined number of 5mdC in the sequence. Two results from the ^13^C NMR data obtained from these oligonucleotides demonstrated the feasibility of this approach (Supplementary Fig. 2). First, we specifically detected the ^13^C signal from the methyl groups on 5mdC in the DNA. Second, we identified the chemical shift of interest for the ^13^C-5mdC (∼15 ppm) for the subsequent brain experiments.

One important challenge for in vivo ^13^C NMR is sensitivity. Even at 100% enrichment, the sensitivity of ^13^C detection is still much lower than ^1^H, which is commonly imaged in vivo. We overcame this challenge by integrating effective labeling using enriched ^13^C-Met, ultrahigh-field imaging systems, and advanced signal processing techniques.

We imaged the ^13^C-labeled brains using a 11.7 Tesla (T) microimaging system (Fig. 3) and obtained whole-sample ^13^C NMR spectra from 8 brains (2 labeled for 10 days, 2 labeled for 32 days, and 2 control unlabeled brains at both 10 and 32 days, Fig. 3C-E). A significant signal difference attributed to ^13^C-5mdC at ∼15 ppm (as expected based on the data from the synthetic oligonucleotides) was observed between the control and labeled brains, and the 32-day labeled brain exhibited significantly stronger signal than the 10-day labeled and control brains (Fig. 3F). The whole-sample spectra were parametrically fitted using an in-house method to quantify the ^13^C NMR 5mdC signal (Supplementary Fig. 3). The signal increase from 10-day to 32-day labeling matches well with the increase in labeling percentage measured by LC-MS/MS analysis and again significantly outpaced brain volume growth (Fig. 3G). These results confirm that increased labeling and the resulting eMRI signals are not driven solely by increases in brain volume and cell density, and dynamically responsive to neurodevelopmental changes.

**Fig. 3.**
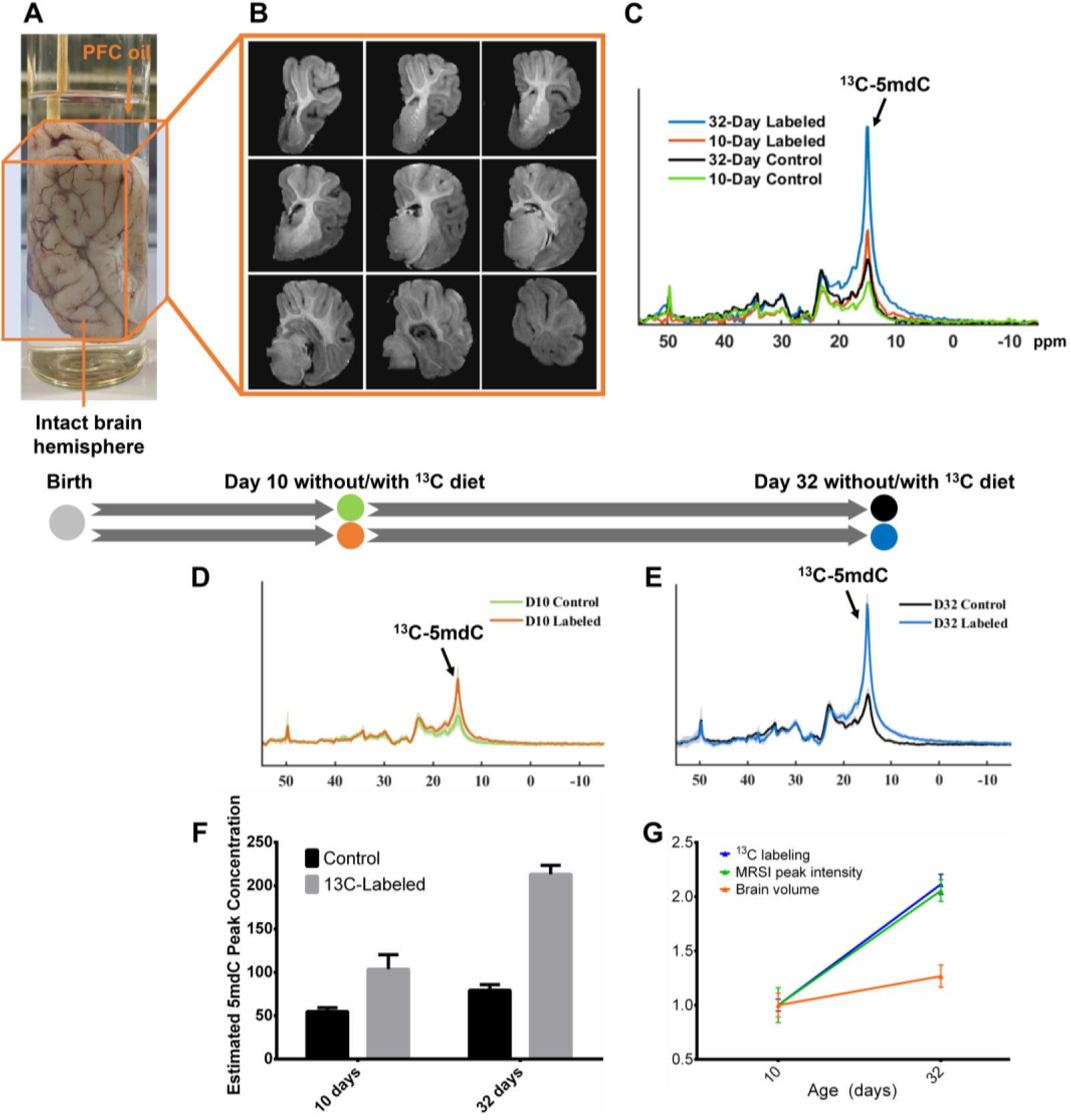
^13^C NMR detects labeled DNA and age-dependent labeling in the brain. **A**, Sample setup for spectroscopy and imaging experiments. The intact brain hemisphere was submerged in PFC oil for susceptibility matching to improve magnetic field homogeneity. **B**, A 3D MR image from one of the brain samples. **C**, Whole-sample ^13^C NMR spectra acquired from different brains at different time points and dietary conditions, i.e., 10-day control diet (green), 10-day ^13^C-Met diet (orange), 32-day control diet (black), and 32-day ^13^C-Met diet (blue). Data for each sample were acquired in 9 h experiments. **D** and **E**, Eight brain hemispheres were measured, four for each time point and two for each dietary condition among the four (without and with ^13^C-Met diet). Consistent results were obtained from each group. Labeled samples, D10 (10 days) in panel D or D32 (32 days) in panel E, produced significantly stronger ^13^C-5mdC signals (at ∼15 ppm, black arrows) compared to age-matched controls. **F**, Brains labeled for 32 days exhibited significantly stronger signals than those labeled for 10 days. The signal increase for the controls from 10 to 32 days was due to brain growth and is attributed to natural-abundance ^13^C NMR signals from 5mdC and methyl groups on thymidine, with the latter a larger contributor due to higher abundance. Nevertheless, the 10-day labeled sample still showed a stronger signal than the 32-day control, indicating the signals measured are primarily from ^13^C-5mdC labeling. **G**, The signal increase from 10-day to 32-day labeled brains approximately matches the labeling percentage increment measured by LC-MS/MS (Fig. 2F) and substantially outpaced brain volume growth.

### Regional differences in brain DNA methylation revealed by eMRI

The ability of ^13^C NMR spectroscopy to noninvasively detect ^13^C-5mdC signals from genomic DNA establishes the premise of using ^13^C-MRSI to image brain DNA methylation in vivo by mapping ^13^C-labeled 5mdC, another key feature of eMRI. We tested this possibility by performing ^13^C-MRSI with a novel subspace imaging strategy (Methods) on brain samples to image DNA methylation.

Our ^13^C-MRSI method robustly mapped the spatial distribution of ^13^C-5mdC in brain. The 32-day labeled brains produced clearly discernible regional variation in ^13^C-5mdC levels (Fig. 4A, B, Supplementary Fig. 4). The localized ^13^C NMR spectra extracted from several representative anatomical brain regions confirmed the regional signal variations attributed to the ^13^C-5mdC peak at ∼15 ppm (Fig. 4C). The putamen, caudate, and thalamus regions showed the strongest signals, while the temporal and occipital lobes were among the regions showing weaker signals (Fig. 4C, D). These regional measurements were confirmed to yield strong and statistically significant differences using pairwise Kruskal-Wallis with Dunn’s multiple comparison tests (Fig. 4E). To validate our ^13^C-MRSI method and the regional variation observed, we performed phantom experiments using NMR tubes filled with ^13^C-5mdC solutions of different concentrations. Concentration estimates by our method were in excellent agreement with the true concentrations (r > 0.9, Supplementary Fig. 5), confirming the accuracy of our ^13^C-MRSI method.

**Fig. 4.**
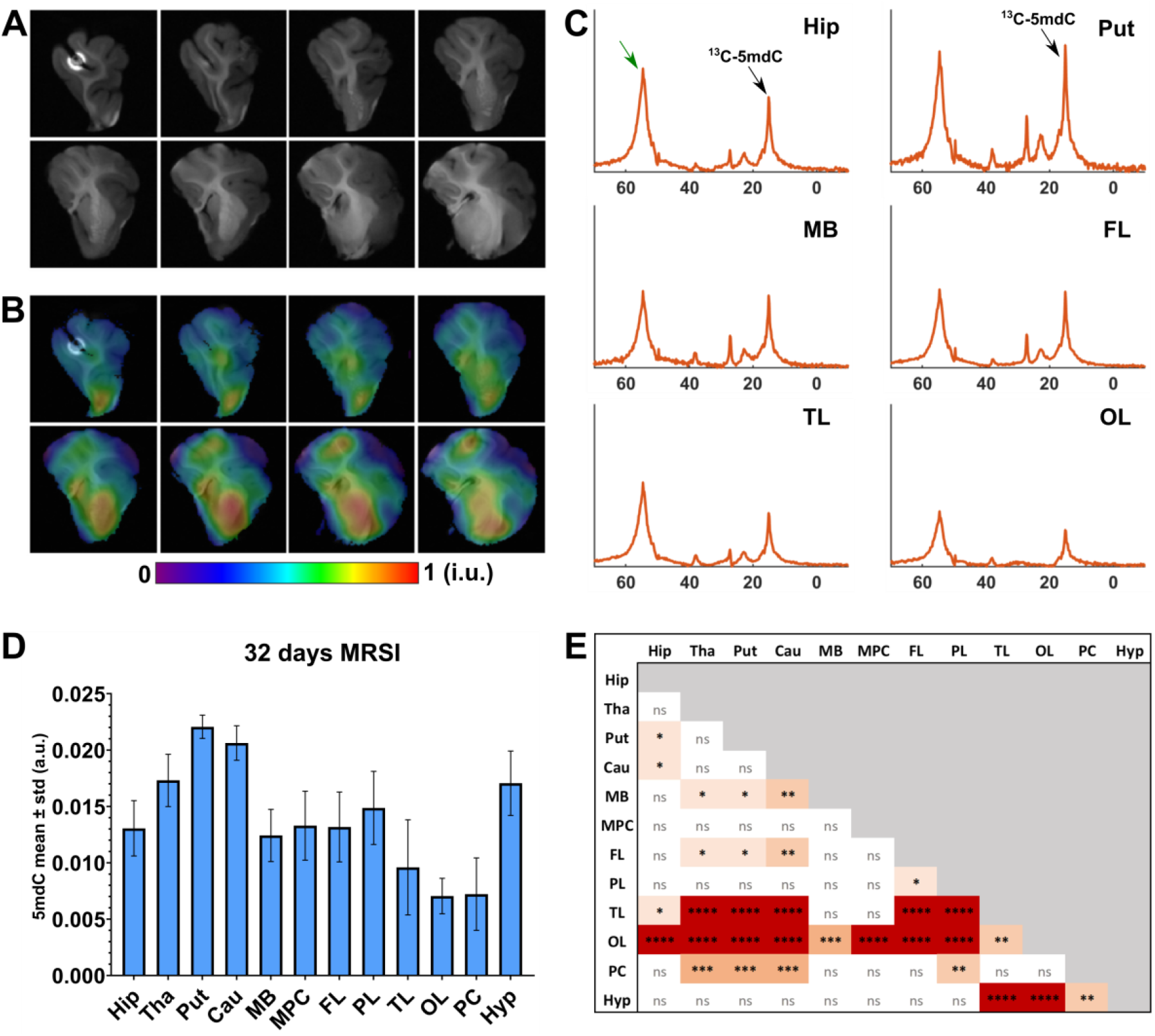
Mapping regional variation of ^13^C-labeled DNA in the intact brain. **A**, Anatomical MR images corresponding to different slices across the intact 3D imaging volume. **B**, Spatial maps of labeled DNA methylation overlaid on the same images in A. The maps were normalized with a maximum intensity of 1 (arbitrary units). Clear spatial variation of labeled DNA methylation is observed. **C**, Localized ^13^C NMR spectra from different brain regions (averaged for the voxels within the same region) also exhibited clear regional variation for the peak assigned to ^13^C-5mdC. The peak at ∼55 ppm indicated by the green arrow is hypothesized to be related to lipid metabolism (Supplementary Fig. 3). **D**, Quantitative comparison of ^13^C-5mdC signal differences across brain regions (error bars indicate the standard deviations for individual regions). Cerebellum and parts of brain stem were not included because they did not fit into the RF probes used in the imaging system. **E**, Pairwise Kruskal-Wallis with Dunn’s multiple comparison tests reveal that several regions produced significantly different ^13^C-5mdC signals and thus different amounts of labeled methylated DNA. * P < 0.05, ** P < 0.001, *** P < 0.0002, **** P < 0.0001. We imaged half brain due to size limitations in our imaging system with the available ^13^C coil. However, this is not an inherent limitation of eMRI, and imaging the whole brain will offer stronger signals. Brain regions not specified in Fig. 2: putamen (Put), caudate (Cau), frontal lobe (FL), parietal lobe (PL), temporal lobe (TL), occipital lobe (OL), hypothalamus (Hyp).

Regional differences in ^13^C-5mdC contents measured by eMRI were much stronger compared to the labeling percentage by LC-MS/MS (Fig. 4D vs. Fig. 2E). We propose that eMRI is providing more sensitive representations of the regional differences in total 5mdC content, thus global DNA methylation. This is supported by global methylation levels measured in tissue samples from selected regions with the most significant eMRI signal differences (i.e., putamen and temporal lobe), using an enzyme-linked immunosorbent assay (ELISA). Higher global methylation was measured in putamen as compared to temporal lobe (Supplementary Fig. 6), consistent with what was observed by eMRI, supporting our hypothesis. Reducing the acquisition time by a factor of 5 still yielded consistent regional ^13^C-5mdC level estimates as compared to the original longer scan (Supplementary Fig. 7), supporting the robustness of eMRI in providing region-specific methylation measures.

## Discussion

^13^C-Met is incorporated in metabolic pathways that lead to the methylation of various molecules in the brain in addition to DNA, e.g., RNA and proteins, potentially generating confounding signals in the ^13^C-MRSI. However, the methyl and other alkyl groups on natural protein side chains have fairly large ^13^C NMR chemical shift differences from the methyl group on 5mdC and should not contribute substantially to what we observe at ∼15 ppm. While various types of methylated RNA may be present in the brain, most methyl groups on these molecules have sufficiently different chemical shifts from 5mdC in the DNA, except for methylated cytosines in the RNA (5mC). The level of mRNA 5mC is estimated at ∼0.03–0.1% of cytosines, which is much rarer compared to ∼4–5% of methylated cytosines in DNA ^42^. Higher methylation levels in rRNA and tRNA have been reported in mouse embryonic stem cells ^43^, but the RNA 5mC levels in mammalian brains remain largely unknown. Furthermore, the turnover rate for RNA is more rapid than for DNA ^44^. Therefore, we anticipate that the contribution from RNA 5mC should not significantly affect the regional variation observed by eMRI (Supplementary Fig. 7).

Another potential signal source to consider is the methyl groups on thymidine (T) in DNA, which have a very similar chemical shift as 5mdC (∼1–2 ppm difference; Supplementary Fig. 1). However, because T is not labeled by ^13^C-Met, the confounding background signals should predominantly come from natural-abundance ^13^C in T. Such signals are weak compared to those from labeling, especially for the 32-day labeled samples (Fig. 3E). For future samples with lower labeling percentage, this background signal can be computationally removed by performing a baseline scan before labeling followed by subtraction.

Regional differences in brain DNA methylation have been reported for frozen samples of human brain tissue ^45^ but cannot be measured in intact brains due to technology limitation. eMRI bridges this gap. Our analysis of the Allen Human Brain Atlas (ABA) reveals regional differences in global gene expression levels (Supplementary Fig. 8), which also vary with age, highlighting the importance of an in vivo approach to measure these differences. The regional gene expression levels from the ABA do not match the regional DNA methylation levels from the current eMRI data in any simple way, including the well-established negative relationship between gene expression and DNA promoter methylation (Supplementary Fig. 8). One possible reason is that the ABA data were from human adults while our eMRI data are from young pigs. In addition, the canonical inverse relationship between DNA methylation and gene expression exists mainly for methylation in gene promoters, whereas eMRI measures total DNA methylation. An additional possible contributing factor to the regional eMRI signal variation is regional differences in DNA density, i.e., quantity of DNA per unit mass. While nonuniform cell densities in different mammalian brain regions have been reported ^46^, it is presently unclear how this nonuniformity translates to DNA density distributions. These results point to the need for more quantitative experiments on gene expression and DNA methylation, to better understand the eMRI signal.

In considering potential applications, it is important to note that eMRI is able to provide information on the turnover of DNA methylation in the brain in vivo. In piglets, which have growing brains, the ^13^C label can be incorporated into the DNA of brain cells through both new cell formation and turnover of DNA methylation, thus increasing the resulting ^13^C labels available for MRSI detection. In adult animals and humans, however, brain growth is not a major factor, so incorporation of ^13^C labels into the DNA of adult brain cells is primarily through turnover of DNA methylation ^47^. No previously known in vivo technique can measure this turnover, and doing so with eMRI may shed light on how the dynamics of brain DNA methylation contribute to the regulation of higher brain functions.

An important consideration for applying eMRI to humans is sensitivity. Dietary administration of ^13^C-Met to adult humans is expected to yield a lower labeling percentage in the brain than for developing piglets ^33^. However, human brains have ∼10× larger volume than the piglet brains imaged here, thus providing significantly larger quantities of 5mdC for signal detection. It is not difficult for humans to consume a Met deficient diet and take ^13^C-Met as a daily supplement.

Since there is evidence supporting that the half-lives of DNA methylation events are measured in hours^47^, it likely will take less than 10 days to reach maximum labeling in humans, making this technology easily translatable. Imaging in ultrahigh-field human MRI systems (7 T or higher) should also provide a significant sensitivity boost. Furthermore, many potential optimizations in data acquisition and processing can be performed for enhancing the ^13^C-MRSI signals.

Although DNA methylation influences gene expression on a per-gene basis, mapping global brain DNA methylation likely provides a general measure of gene expression activity. This is analogous to how fMRI measures general neuronal activity without knowledge of specific neurons. There is growing evidence suggesting the functional significance of global methylation changes, e.g., in psychiatric disorders and cancer^48,49^. eMRI may enable in vivo epigenetic study at long time scales from days to months, where such time scales are experimentally inaccessible by radioactive labeling and invasive techniques. eMRI can thus be coupled with fMRI to investigate the interaction between short-term neural and long-term molecular control of brain function, to gain further insights into the regulation of behavior and brain responses to environmental and disease stimuli.

## Materials and Methods

### DNA synthesis

DNA oligonucleotides were prepared by solid-phase synthesis on an ABI 394 instrument using reagents from Glen Research, including the 5-methyl-2′-deoxycytidine (5mdC) phosphoramidite. Oligonucleotides were purified by 7 M urea denaturing 20% PAGE with running buffer 1× TBE (89 mM each Tris and boric acid and 2 mM EDTA, pH 8.3), extracted from the polyacrylamide with TEN buffer (10 mM Tris, pH 8.0, 1 mM EDTA, 300 mM NaCl), and precipitated with ethanol. The 20-nucleotide DNA sequence, designed to avoid self-dimer formation and to include C nucleotides in various sequence contexts, was 5′-CTACGCCTCGCTCGCCCCTT-3′, where the ten C nucleotides were either all 5mdC or all standard dC (MW 6,081 with 5mdC). Approximately 750 nmol of each oligonucleotide was dissolved in 0.5 mL 10% D_2_O/90% H_2_O (1.5 mM oligonucleotide; 9 mg/mL) in a standard 5 mm NMR tube for ^1^H and ^13^C NMR spectroscopy.

### ^13^C-MRSI phantom

^13^C-labeled 5mdC (5-Methyl Cytosine-^13^C, ^15^N_2_ Hydrochloride 99%, Cat. M294702, from Toronto Research Chemicals) was dissolved in PBS (Gibco) then serially diluted to the desired concentrations of 1 mM, 4 mM, 6 mM and 10 mM. The solutions were transferred to 10 mm SP Scienceware Thin Walled Precision NMR tubes (Wilmad-LabGlass) which were taped together to fit into the 30-mm probe of the 11.7 T microimaging system.

### Piglet experiments and dietary labeling

All animal care and experimental procedures were in accordance with the National Research Council Guide for the Care and Use of Laboratory Animals and were approved by the University of Illinois Institutional Animal Care and Use Committee (Protocol #20223). Naturally farrowed, intact domestic male pigs derived from the cross of Line 2 boars and Line 3 sows (Pig Improvement Company) were obtained from a commercial swine herd and transferred to the Piglet Nutrition and Cognition Laboratory (PNCL) on post-natal day (PND) 2 (n=8). Pigs were maintained at PNCL through PND 32 per standard protocols as described elsewhere with the following modifications. All pigs were provided *ad libitum* access to water and a custom, nutritionally complete ^50^ milk replacer formula (TestDiet) via a semi-automated liquid delivery system that dispensed milk from 1000 to 0600 h the next day ^51^. The milk replacer powder was formulated to contain 30.4% lactose, 30.3% stabilized lipids (i.e., dried fat; combination of lipid sources including sunflower oil, palm oil, and medium-chain triglycerides), and 25.4% purified amino acids; no intact protein was included in this formulation. The nutritional profile of the milk replacer powder included 0.50% methionine and 0.49% cysteine, with the control and labeling diets containing regular L-methionine (Met) and *methyl-*^13^C-L-methionine (^13^C-Met, 99%, Sigma-Aldrich), respectively (i.e., all Met present in the labeling diet contained the stable ^13^C isotope label). Milk replacer formula was reconstituted fresh daily with 200 g of milk replacer powder per 800 g water. Pigs were individually housed in caging units (87.6 cm × 88.9 cm × 50.8 cm; L × W × H), which allowed each pig to see, hear, and smell, but not directly touch, adjacent pigs. A toy was provided to each pig for enrichment, and pigs were allowed approximately 15 min of direct social interaction with other pigs once daily. The ambient environment in the rearing space was maintained on a 12 h light and dark cycle from 0800 to 2000 h, with room temperature set at 27 °C for the first 21 days of the study and gradually lowered to 22 °C during the last 7 days.

### Brain biopsy and fixation

At the appropriate age (PND 10 and 32 days), each pig was humanely euthanized per standard protocol ^52^ and the whole brain was quickly excised post-mortem or when meeting euthanasia criteria ^53^. The brain was separated into two hemispheres, with tissue samples collected from 8 different regions within the left hemisphere (medial prefrontal cortex, cerebellum, hippocampus, midbrain, brain stem, thalamus, perirhinal cortex, and striatum). Enough tissue to permit the ultimate extraction of at least 1 μg total DNA (approximately 100-250 mg) was snap-frozen on dry ice and stored at –80 °C for genomic DNA extraction. The right hemisphere from each brain remained intact (i.e., was not biopsied), and was fixed in 10% neutral-buffered formalin (NBF) at 4 °C for 72 h and stored in PBS (pH 7.4; Gibco) at 4 °C for subsequent NMR spectroscopy and imaging experiments.

### Genomic DNA extraction

Pulverized tissue samples were incubated in a digestion mix (100 mM Tris, 5 mM EDTA, 200 mM NaCl, 0.2% SDS, pH 5.5) containing 400 μg ml^-1^ Proteinase K (Invitrogen) and 200 ug ml^-1^ RNase A (Invitrogen) at 50 ºC overnight. Genomic DNA was extracted with phenol:chloroform:isoamyl alcohol (25:24:1, Fisher Scientific) and precipitated using 70% ethanol (Sigma-Aldrich). Genomic DNA was resuspended in HPLC-grade water (Sigma-Aldrich) and the concentration measured with the NanoDrop Spectrophotometer (Thermo Fisher Scientific).

### LC-MS/MS DNA analysis

1 μg of genomic DNA was hydrolyzed overnight at 37 ºC with 5 U of DNA Degradase Plus (Zymo Research) following the manufacturer’s protocol in 50 μl volume. Digested DNA was filtered through Amicon Ultra 10K centrifuge filters (EMD Millipore) to remove undigested polynucleotides and collected for LC-MS/MS. LC-MS/MS analysis of ^13^C-5mdC isotope labeling was performed with a Waters SYNAPT G2Si mass spectrometer and a Waters ACQUITY UPLC H-class fitted with a Waters Cortecs C18+ column (2.1 × 150 mm, 1.6 μm particle size) at the flow rate of 150 μl min^-1^ under a gradient of ammonium acetate and acetonitrile (Fisher Scientific). Multiple reaction monitoring (MRM) was set up to capture the transitions from deoxycytidine – cytosine (228.1-112.05 Da), 5-methyldeoxycytidine to 5-methylcytosine (242.1-126.07 Da), and ^13^C-5-methyldeoxycytidine to ^13^C-5-methylcytosine (243.1-127.07 Da). The percentage of ^13^C isotope labeling was determined by the following formula using peak areas of dC and 5mdC:

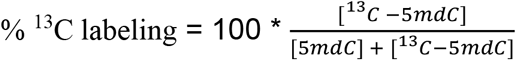

### Global DNA methylation by ELISA

Brain tissues were collected from pigs at 32 days of age (n=4; 1 female, 3 males). Putamen and temporal lobe samples (∼25 mg each) were dissected from the left hemisphere of each pig and briefly homogenized with a scalpel. Samples were then flash-frozen in liquid nitrogen and stored in cryovials at –80 °C. Genomic DNA was extracted using the DNeasy Blood & Tissue Kit (Qiagen). A Nanodrop 2000 (Thermo Fisher Scientific) was used to determine DNA concentrations/yield. As a quality control check, all A_260_/A_280_ ratio values indicated sufficient DNA purity (values ranged from 1.87 to 2.02). Global DNA methylation was assessed via an ELISA-based commercial kit [MethylFlash Methylated DNA Quantification Kit (Fluorometric); Epigentek]. All standards and samples were run in duplicate. Absorbance was read at 450 nm using a plate reader (BioTek) and Gen5 version 3.11 software. DNA methylation levels in each brain region were calculated relative to the Methylated Control DNA standard using the single point method equation: [(Average A_450_ Sample – Average A_450_ Blank) / ((Average A_450_ Methylated Control DNA – Average A_450_ Blank) x 2)] x 100%. For calculations, the quantity of each unknown sample DNA matched the quantity of Methylated Control DNA (i.e., the blanked A_450_ value for a 100 ng DNA sample was compared to the blanked A_450_ value for 100 ng Methylated Control DNA), and the factor of 2 was to normalize the percentage of 5mC in the control DNA to 100% as suggested in the manufacturer’s instructions. This calculation yields relative global methylation level with respect to the control DNA, which is sufficient for our intended regional comparison.

### Statistical analysis

One-way ANOVA followed by Tukey’s multiple comparison test was used to compare the ^13^C labeling efficiencies between different brain regions as measured by LC-MS/MS. Kruskal-Wallis followed by Dunn’s multiple comparison tests were used to compare the ^13^C-MRSI signal intensities from the different brain regions of the 32-day labeled brain.

### Brain NMR spectroscopy and imaging experiments

Each intact piglet brain hemisphere was placed in a 30 mm diameter glass tube containing perfluorocarbon oil (Fluorinert FC-40, Sigma-Aldrich) as a magnetic susceptibility matching fluid. A wooden applicator stick was taped in place to hold the brain tissue and prevent it from floating. All brain NMR spectroscopy and imaging experiments were performed using a Bruker AV3HD 11.7 Tesla/89 mm vertical-bore microimaging system equipped with a 16-channel shim insert and Micro2.5 gradient set with a maximum gradient of 1500 mT/m. Data were collected using a 30 mm dual-tuned ^1^H/^13^C RF resonator and ParaVision 6.0.1 (Bruker Biospin). A 200 mM ^13^C-Met solution dissolved in PBS was used for ^13^C calibration before the brain imaging scans.

### MRI

During each experiment session, a localizer scan was first acquired, followed by field map shimming. The B0 inhomogeneity was minimized to reach a ^1^H water linewidth of between 45 and 60 Hz. Anatomical MRI scans were acquired using axial RARE (T2 weighted) and MDEFT (T1 weighted) with the following parameters, RARE: TR/TE = 6000/5.5 ms, RARE factor = 4, number of averages (NA) = 2, field of view (FOV) = 30 × 30 mm, matrix size = 192 × 192, and 50 1 mm slices; 3D MDEFT: TR/TE = 4000/2.2 ms, inversion delay = 1050 ms, FOV = 30(x) × 30(y) × 50(z) mm^3^, matrix size = 192 × 192 × 50, and 8 segments. A field map was collected covering the same FOV. Brain volume was estimated from the T1-weighted MR image for each sample. The MR image was also segmented into 12 regions of interest (ROIs), i.e., hippocampus, thalamus, putamen, caudate, midbrain, medial prefrontal cortex, frontal lobe, parietal lobe, temporal lobe, occipital lobe, perirhinal cortex and hypothalamus, for regional analysis of MRSI results. The regions for tissue sampling (for LC-MS/MS and Fig. 2) were defined slightly differently from those for MRI-based segmentation, due to the difficulty in precisely extracting some regions from the piglet brains (e.g., putamen and caudate were assigned as striatum in the LC-MS/MS analysis). The cerebellum and brain stem were excluded in the imaging experiments because of the size limitation of the instrumentation used.

### ^13^C NMR spectroscopy and MRSI

A whole-sample ^13^C NMR spectrum was collected at 125.755 MHz using a 100 μs RF pulse centered at 20 ppm, with a 2 sec TR. Broadband ^1^H decoupling was achieved with a WALTZ-16 decoupling scheme centered at 2 ppm (^1^H carrier frequency). The free induction decay (FID) was collected with a 22727 Hz (∼180 ppm) spectral bandwidth (BW), 1024 points, and 16238 averages for a total acquisition time of 9 h and 6 min. ^13^C-MRSI data was acquired using a phase-encoded chemical shift imaging (CSI) sequence with a 50 μs, 30°, block RF excitation pulse, a 30 × 30 × 50 mm^3^ FOV, an 8 × 8 × 8 matrix, and broadband ^1^H decoupling described above. FIDs were collected with the same BW and number of points, TR/TE = 2000/1.16 ms and 64 averages for a total acquisition time of 18 h and 12 min. A matching ^1^H-CSI data was collected with TR/TE of 500/1.16 ms, 16 × 16 × 8 matrix size, and 512 FID points over a 20 ppm BW (17 min), to correct for spatial misalignment between the MRI and MRSI acquisitions.

### NMR and MRSI data processing

A model-based data processing method was developed to perform spectral quantification of the measured ^13^C-MRSI data to obtain the ^13^C-5mdC concentration maps for eMRI. The method incorporates both physics-based prior information (resonance structure of the ^13^C NMR spectrum) and the high SNR single-voxel spectroscopy (SVS) data from the whole sample. We first performed spectral quantification of the high SNR SVS data, *sSV*(*t*), to derive spectral basis functions for our model. We expressed *sSV*(*t*) as:

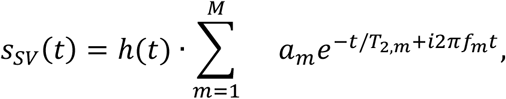

where *am, fm*, and *T*_2,*m*_ represent concentration, resonance (chemical shift) frequency, and spin-spin relaxation constant of the *m*th spectral component, respectively, and *h*(*t*) accounts for non-Lorentzian spectral lineshape variations introduced by non-ideal experimental conditions (e.g., magnetic field drift, field inhomogeneity, etc.). In this work, *h*(*t*) corresponds to a compensated Gaussian lineshape function so that the spectral model can be fit to the experimental data, with the fitting residual at the noise level and passing the Komolgrov Gaussianity test (Supplementary Fig. 3).

After spectral quantification of the SVS data, we obtained the following spectral basis functions that contain resonance frequencies and spin-spin relaxation constants of all spectral components and the spectral lineshape compensating function:

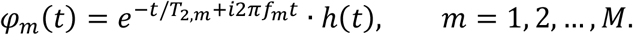

To perform spectral quantification of the entire ^13^C-MRSI data set, we first mapped the data to the spatial domain using a constrained image reconstruction method that includes B_0_ field inhomogeneity correction ^54^. This reconstruction step included some spatial filtering effects to enhance SNR of the reconstruction. The spectral basis functions derived from the whole-sample SVS data above were used to quantify the reconstructed spatiotemporal function *ρ*(*x, t*) point by point. Considering the difference in spatial resolution, shimming condition, and sequence set up between the SVS and MRSI scans, we adjusted the spectral basis functions *φ*_*m*_(*t*) learned from the SVS data to better match the MRSI data. To this end, we first denoised the reconstructed spatiotemporal function *ρ*(*x, t*) using a low-rank filtering method ^55^. We then estimated a new spectral lineshape function *h*(*t*) from the denoised MRSI data. The estimated lineshape function was applied to *φ*_*m*_(*t*) to obtain an improved set of spectral basis, {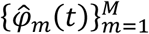. With 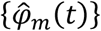} determined, concentrations, *ĉ*_*m*_ (*x*), of all spectral components were determined from the original reconstructed spatiotemporal function *ρ*(*x, t*) data by solving the following model fitting problem:

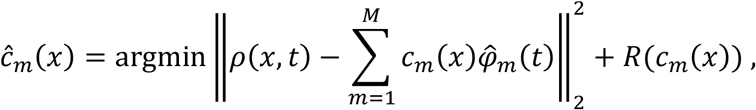

where *R*(·) represents an edge-weighting regularization function to incorporate spatial prior information. The spatial distribution of the spectral component corresponding to ^13^C-5mdC was separated. Regional concentrations and spectra were obtained from the 12 ROIs segmented from MRI and analyzed.

## Data and materials availability

All data generated will be made available upon reasonable request to G.E.R. and K.C.L.. Custom codes were developed to analyze the ^13^C-NMR and ^13^C-MRSI data. The codes will be made available upon reasonable request to G.E.R. and K.C.L..

## Acknowledgments

The authors thank Dean Olson and Lingyang Zhu for discussions on ^13^C NMR spectroscopy and acquiring the ^13^C NMR data for the synthesized oligonucleotides; Furong Sun for developing the LC-MS/MS protocol and instruction on mass spectrometry data analysis; Cong Zhou for his help with solid-phase DNA oligonucleotide synthesis; Yibo Zhao for his help with ^13^C MRSI data processing; Adam Jones for his help on piglet feeding and rearing; Joanne Fil for her help on piglet brain tissue dissection; and Katie Ranard for her help on the DNA methylation measurement using ELISA kits. Thanks also to the Beckman Institute for Advanced Science and Technology and the Carl R. Woese Institute for Genomic Biology for access to facilities and technical support.

## Author contributions

F.L. contributed to concept development, technical implementation, data collection, data analysis, and manuscript writing. J.C. contributed to technical implementation, data collection, data analysis, and manuscript writing. J.S.C. contributed to technical implementation, data collection, and data analysis. C.C. contributed to technical implementation, data collection, data analysis and manuscript writing. T.K.H. contributed to data collection, data analysis, and manuscript writing. S.K.S. contributed to concept development, technical implementation, data collection, data analysis, and manuscript writing. Z.P.L. contributed to concept development, technical implementation, data analysis, and manuscript writing. R.N.D. contributed to technical implementation, data collection, data analysis, and manuscript writing. G.E.R. contributed to concept development, technical implementation, data analysis, and manuscript writing. K.C.L. contributed to concept development, technical implementation, data analysis, and manuscript writing.

## Supplementary Figures

**Supplementary Fig. 1.**
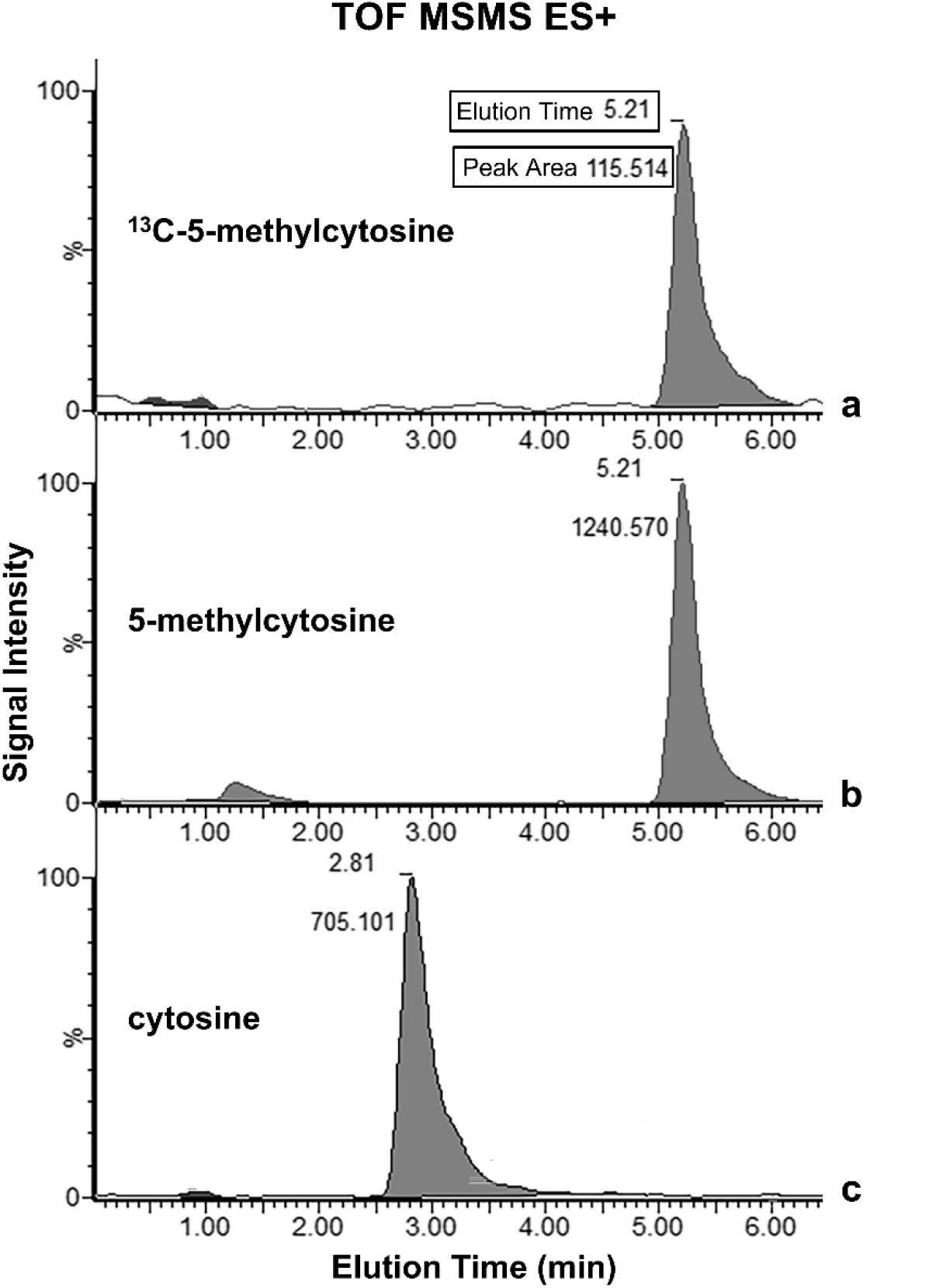
LC-MS/MS data reveal ^13^C-5mdC and 5mdC for quantifying labeling percentage. Representative time of flight (TOF) MS/MS multiple reaction monitor (MRM) data of **A**, ^13^C-5-methylcytosine (fragmented from ^13^C-5mdC after the first MS), **B**, 5-methylcytosine (fragmented from 5mdC), and **C**, cytosine (fragmented from dC). The data here were from genomic DNA isolated from the thalamus of a 10 days ^13^C-labeled piglet. Characteristic elution times and specific molecular weights of the primary ions confirm the identities of the peaks. Area under the peaks were calculated to estimate the ^13^C-enrichment of genomic 5mdC. Elution time are in minutes and peak area are indicated for each peak. 100% signal intensity is set to the intensity of the most prominent peak of the queried molecular weight.

**Supplementary Fig. 2.**
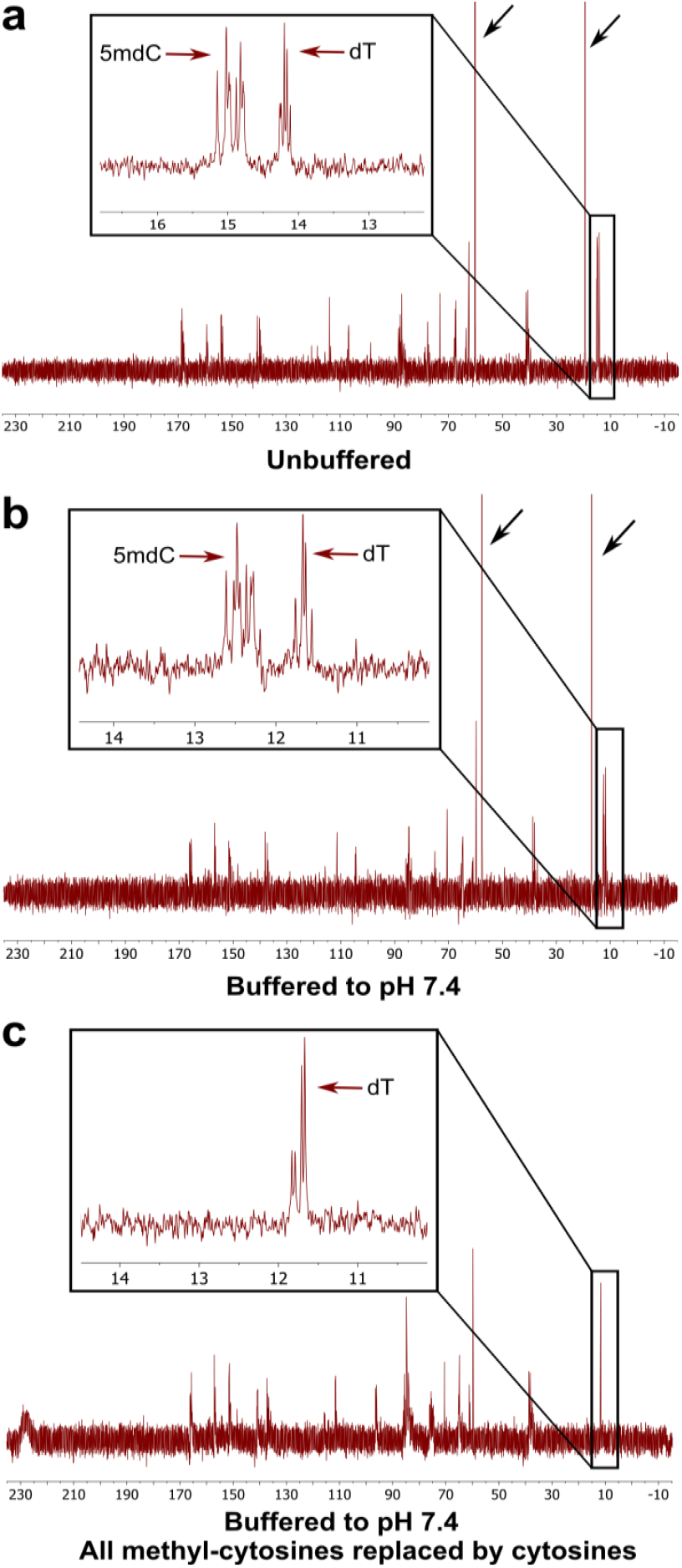
^13^C NMR spectroscopy specifically detects the ^13^C signals from the 5mdC within a synthetic DNA oligonucleotide. (sequence: 5′-CTACGCCTCGCTCGCCCCTT-3′). **A**, High-resolution ^13^C NMR spectrum of the unbuffered sample, acquired with a Bruker Carver B500 NMR spectrometer, 500 MHz (∼11.75 T), equipped with a CryoProbe. The 10 slightly dispersed peaks at ∼15 ppm from the methyl groups on the 5mdC nucleotides at different positions of the sequence are clearly identified and differentiated from the 5 peaks at ∼14.2 ppm from the methyl groups on the dT nucleotides. **B**, The spectrum from the same sample buffered with HEPES to pH 7.4 showed consistent results, with the chemical shift of 5mdC at ∼12.5 ppm. **C**, The spectrum from a control sequence for which all 5mdC were replaced with regular dC; all peaks from 5mdC disappeared, with only peaks from dT remaining. The black arrows in the first two rows indicate peaks from ethanol, which was removed in the last sample. These data provided the information that NMR signals for the ^13^C-labeled 5mdC should be in the range of 12-15 ppm. On this basis, we assigned the increased signal observed at ∼15 ppm in the brain experiments to ^13^C-5mdC.

**Supplementary Fig. 3.**
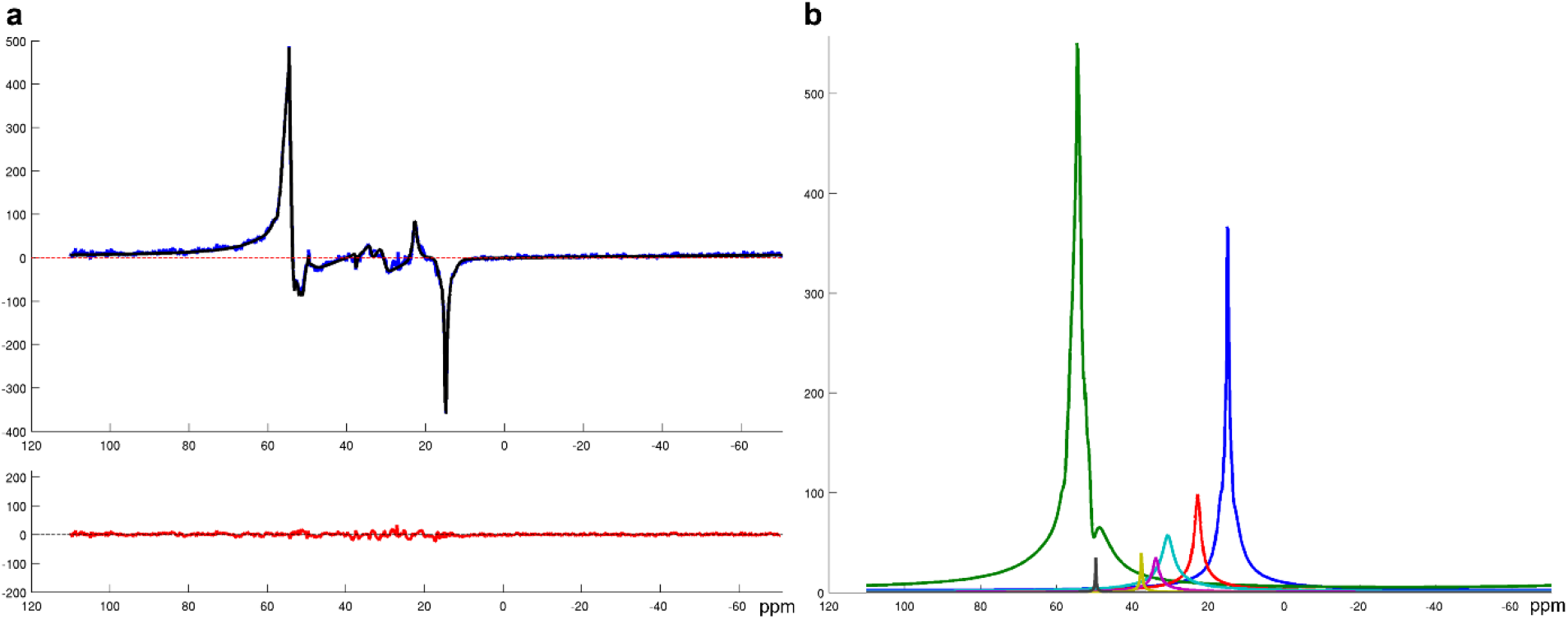
^13^C NMR spectroscopy data and fitting results from a 32-day labeled brain. **A**, High-fidelity fitting of the whole-sample SVS data. The blue curve is the real part of the raw ^13^C NMR spectrum, and the black curve is the model fit. The fitting residual shown below in red is at noise level and passed the Komolgrov Gaussianity test. **B**, Different spectral components extracted from the fitting (coded by different colors and with lineshape distortion *h*(*t*) incorporated). These components were used to construct the basis for fitting the MRSI data. The blue peak (right column) corresponds to ^13^C-5mdC from labeling. The strong green peak centered at ∼55 ppm is hypothesized to be from ^13^C labeled phospholipids through the methylation pathway that converts phosphatidylethanolamine to phosphatidylcholine. This signal is not of interest in the present study and was thus excluded in Fig. 3. But it may be of interest in studies concerning lipid metabolism.

**Supplementary Fig. 4.**
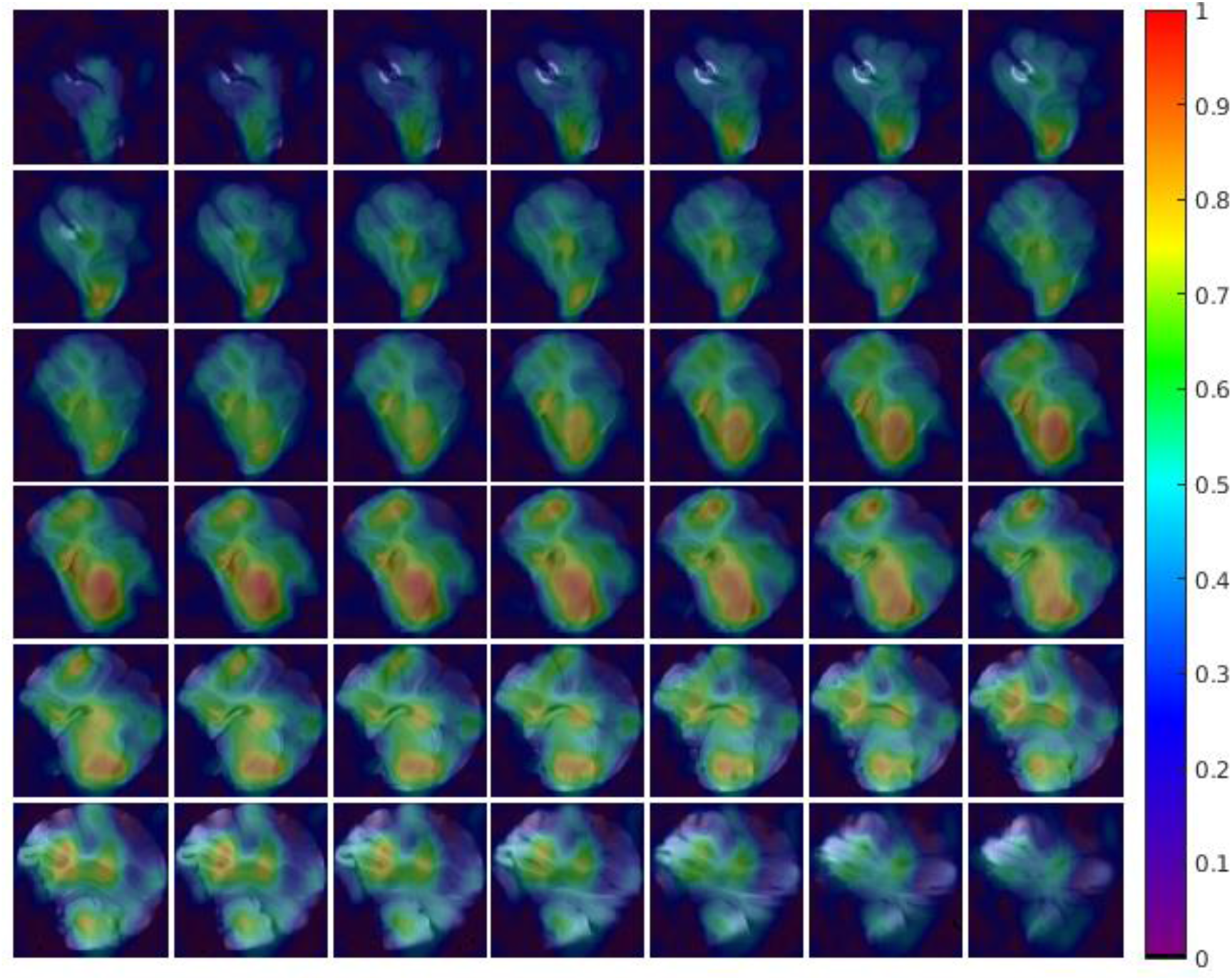
^13^C-5mdC maps across the whole 3D imaging volume. The spatiospectrally reconstructed ^13^C-MRSI data (with a lower resolution than the MR images) from Fig. 4 were first resized to the resolution of the anatomical MR images and then quantified (for better visualization). The quantified ^13^C-5mdC maps were color coded and overlaid on the 3D MRI, normalized to [0,1]. Strong signals can be seen within the brain, with a clear distinction from the nonbrain background noise, implying an excellent signal-to-noise ratio.

**Supplementary Fig. 5.**
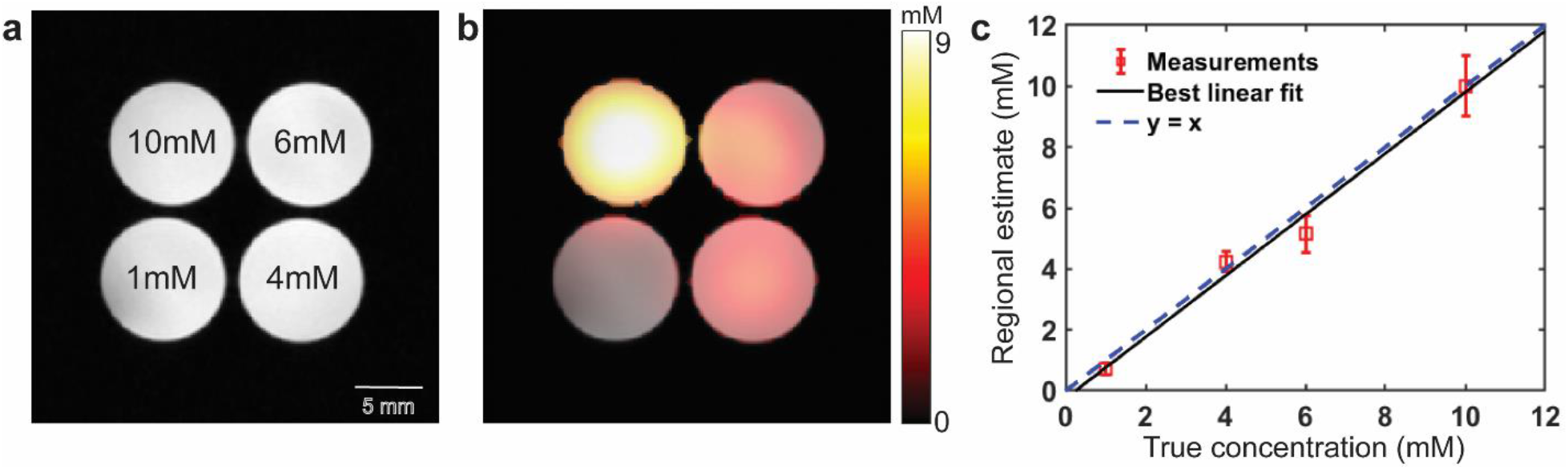
Phantom results for validating our ^13^C-MRSI acquisition and processing methods. **A**, Phantom setup: Four 10 mm NMR tubes were filled with ^13^C-5mdC solutions with varying concentrations and placed in the microimaging system. The concentrations are labeled in the T2-weighted MRI shown. **B**, A ^13^C-5mdC map for one 2D slice from the 3D imaging data, with a spatial resolution of 3.75 × 3.75 × 5 mm^3^. The spatial intensity distribution matches well with the true concentration variations. **C**, Quantitative comparison between regional concentration estimates from individual tubes and the true values. The original estimates were in arbitrary units. The highest value was normalized to 10 mM and regressed against the true values. Accurate results were obtained with a 0.9 correlation coefficient between the estimates and true values. The error bars of the measurements indicate the standard deviations within each region of interest.

**Supplementary Fig. 6.**
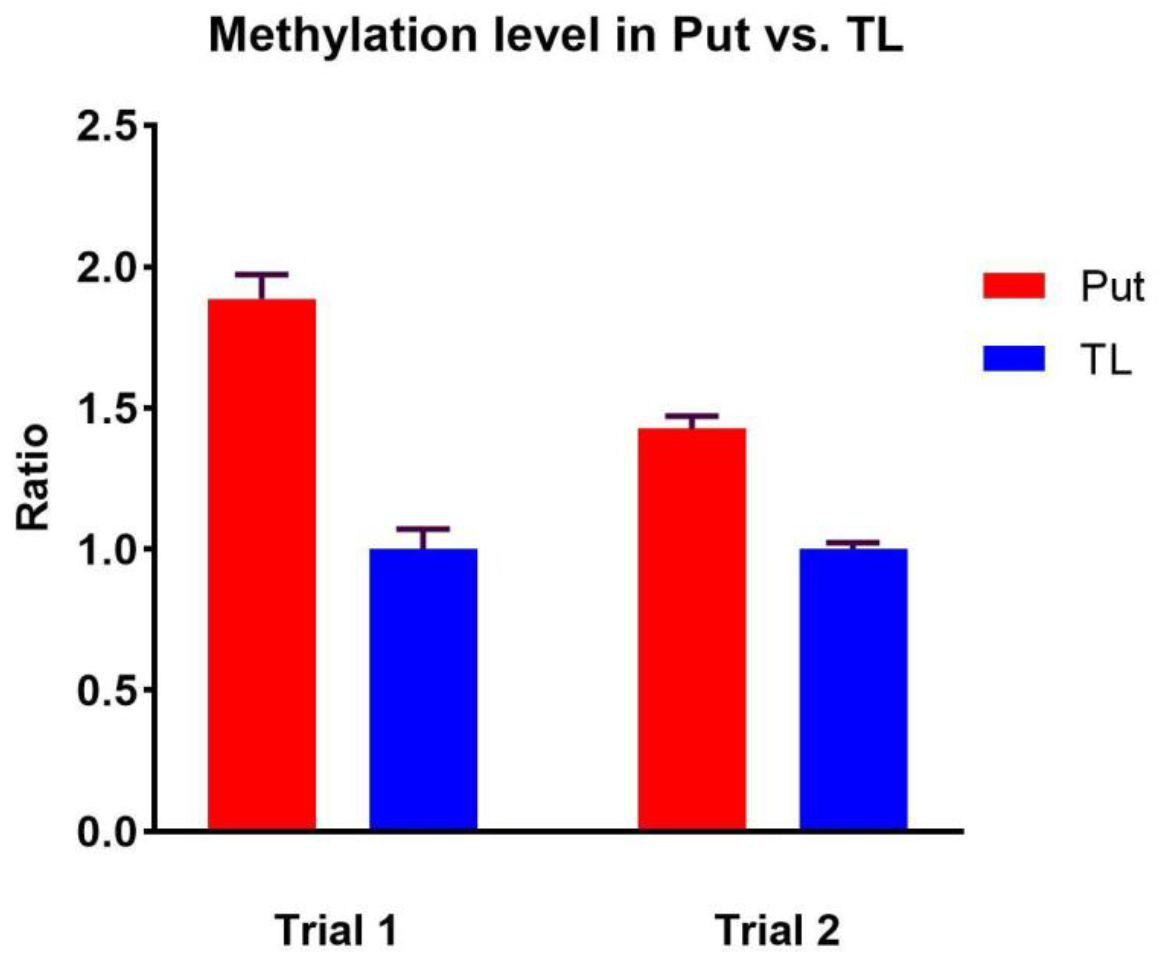
Global DNA methylation in the putamen (Put) and temporal lobe (TL) measured by ELISA is consistent with eMRI data. Relative methylation levels in the Put and TL were measured in two different trials using a fluorometric ELISA kit. Since the goal was to confirm regional differences and the absolute methylation levels were therefore not of interest, we calculated the ratios of methylation levels between Put and TL, with TL methylation level normalized to 1. Both trials showed clearly higher methylation in Put than TL with ratios > 1, consistent with the results from eMRI. Error bars indicate the standard deviations within each brain region. While higher Put methylation was consistently measured, the exact ratios varied between trials 1 to 2 (run in consecutive days) due to fluorescence reading variations.

**Supplementary Fig. 7.**
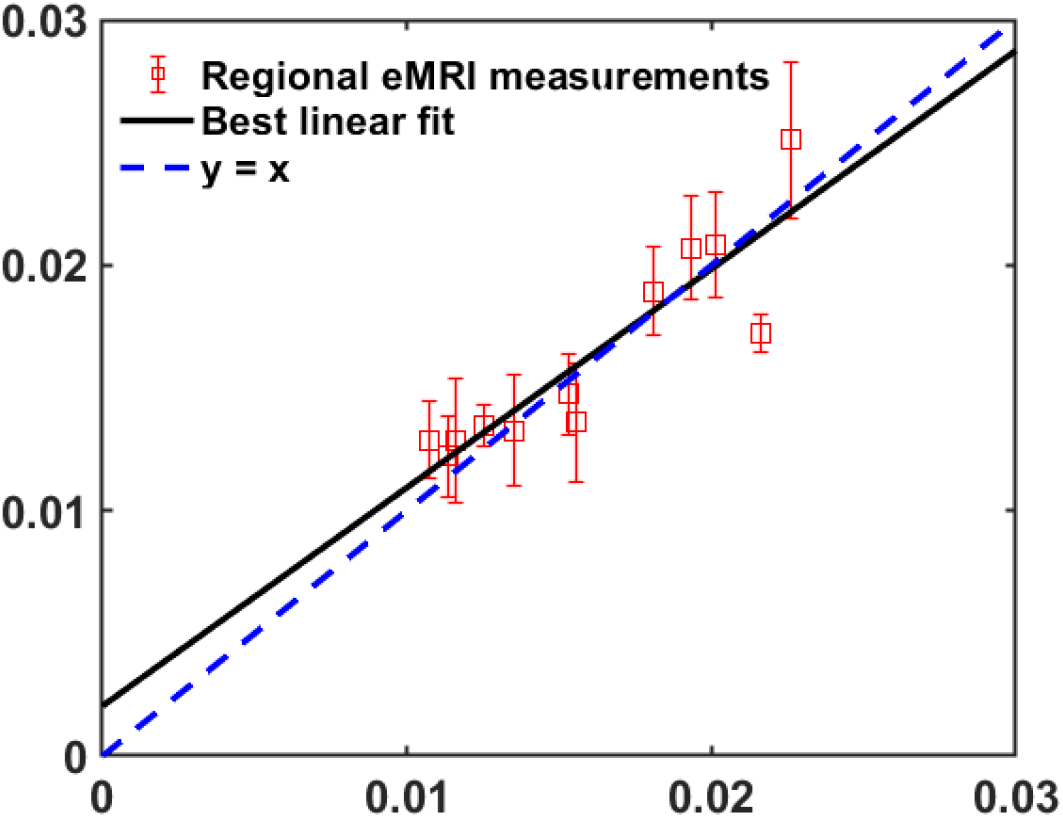
Single repetition ^13^C-MRSI versus five-repetition ^13^C-MRSI (5× longer imaging time). Each repetition in our ^13^C imaging scan took about 18 h. While the mapping results in Fig. 4 were from a five-repetition scan, as shown by the image below, the regional eMRI measurements from a single repetition are consistent with a 5× longer scan (correlation coefficient ∼0.9). This indicates strong translational potential to human experiments. Taking into account the larger brain volume for human, lower labeling efficiency, and the use of a 7 T human system, we predict an approximately 1-2 h scan would be sufficient for high-quality human data. However, since eMRI measures a very stable signal due to the stability of the ^13^C isotope, we have the flexibility to acquire multiple signal averages across several shorter experiment sessions to achieve a sufficient signal-to-noise ratio.

**Supplementary Fig. 8.**
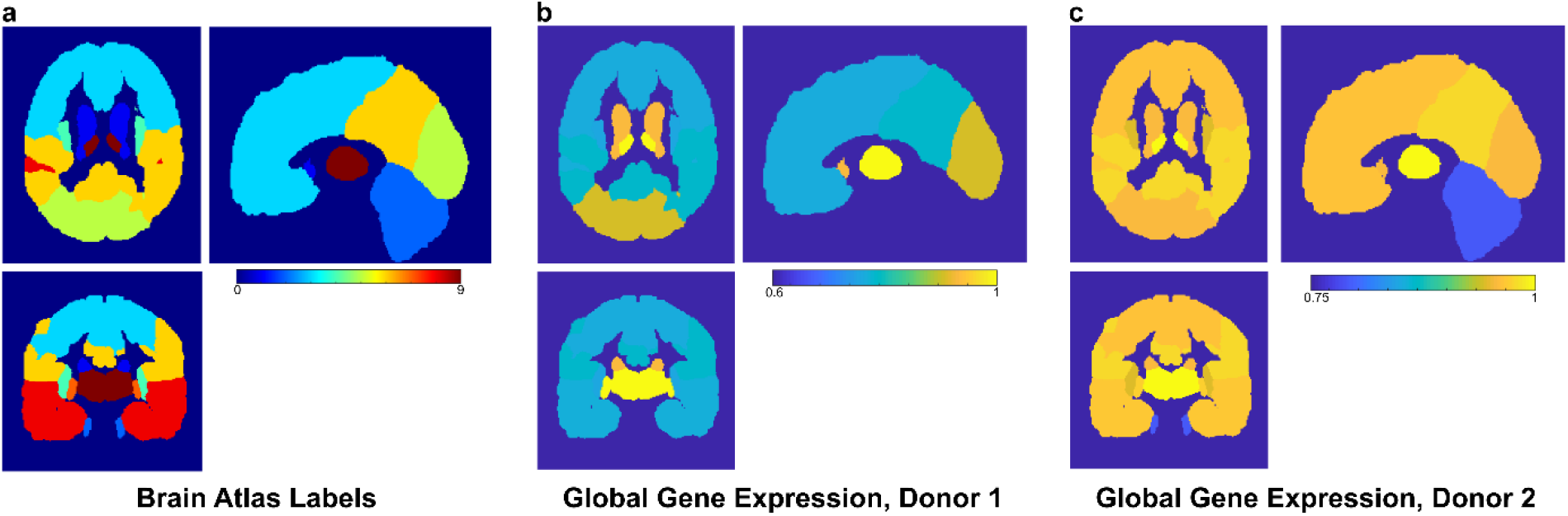
Analysis of the Allen Brain Atlas (ABA) reveals spatial variation in global gene expression in human brains. RNA-sequencing (RNAseq) data from two brains were downloaded from the ABA (https://human.brain-map.org/static/download). The RNASeq data contain transcriptomic profiles for tissues sampled from different brain regions and were used to calculate global gene expression. These global expression values were then mapped to a brain atlas (http://nist.mni.mcgill.ca/mni-average-brain-305-mri/). **A**, A three-plane view of the Montreal Neurological Institute (MNI) brain atlas was shown, with different brain regions segmented from MRI and labeled. Nine anatomical regions were considered: caudate, cerebellum, frontal lobe, insula, occipital lobe, parietal lobe, putamen, temporal lobe, and thalamus. **B**, Maps of global gene expression from one of the two ABA brains. The RNAseq data contain expression data for 22318 genes, each of which has measurements for tissues sampled across different brain structures. We kept only the genes for which the expression levels were reliably detected in all samples. For each brain structure, global gene expression was calculated by summing the expression levels of individual genes, which provided an array of values that were then mapped to the MNI atlas regions in A. Clear regional variation in global gene expression can be visualized. **C**, The same analysis is shown for the second ABA brain. The two brains were from donors at different ages. The global expression values across different structures were normalized using the maximum values for both B and C. Only the brain structures available in both the MNI atlas and RNASeq data were considered. These results support the presence of spatial variation in global gene expression, consistent with our eMRI results. eMRI provides a non-invasive surrogate to explore the functional significance of variation in gene expression for brain function and disease.

